# Discovery of Novel Long Non-Coding RNAs with potential role in zebrafish brain regeneration

**DOI:** 10.1101/2024.06.03.597135

**Authors:** Surbhi Kohli, Dasari Abhilash, Hemlata, Priyanka P. Srivastava, Vishantan Kumar, Shilpi Minocha, Ishaan Gupta

## Abstract

Understanding brain regeneration mechanisms is vital for treating neurological conditions. Zebrafish (*Danio rerio*) are an excellent model due to their genetic similarity to humans and strong regenerative abilities. In this study, we identified novel long non-coding RNAs (lncRNAs) in the regenerating zebrafish brain following traumatic brain injury (TBI). RNA sequencing data of the zebrafish telencephalon from the BioStudies database was analyzed for novel long non-coding RNA expression (lncRNA) at control, one day post-lesion (early wound healing), three days post-lesion (cell proliferation), and 14 days post-lesion (differentiation). We identified 689 potential lncRNAs using HISAT2, StringTie, FEELnc, and PhastCon analysis tools. Principal component analysis (PCA) of identified lncRNAs revealed a distinct expression profile at 1-day post-lesion, indicating their significant role in early wound healing.

Weighted Gene Co-expression Network Analysis (WGCNA) identified two modules (brown and turquoise) showing unique expression patterns critical to brain regeneration. Pathway enrichment analysis linked brown module lncRNAs to peptide biosynthesis, cellular amide metabolism, and ribosome biogenesis. In contrast, turquoise module lncRNAs were associated with ion transmembrane transport and cell adhesion pathways. qPCR validation confirmed co-expression patterns of selected lncRNAs and correlated genes, emphasizing their regulatory roles. This study demonstrates that lncRNAs play crucial roles in zebrafish brain regeneration by modulating gene expression during the early wound healing stage. These insights offer potential therapeutic applications of lncRNAs in neuroregenerative medicine.

## Introduction

In order to effectively treat a variety of neurological conditions and traumas, it is essential to understand the mechanisms behind brain development and injury. With its estimated 100 billion neurons, the human brain is incredibly complex, which presents huge challenges for researchers. For this reason, model organisms that are simple to study but have important structural and functional similarities to the human brain are invaluable.

Zebrafish have shown to be an effective model for researching brain injuries and development. With almost 70% sequence similarity to humans, they are great models for studying brain function, development, and related illnesses. In-depth research on brain morphogenesis can be made possible by the structural parallels between the developing zebrafish brain and the brains of other vertebrates, including humans (Lowery et al. 2009). Zebrafish models have been very helpful in studying neurogenesis from embryonic stages to adulthood. They have shown us how new neurons are created and integrated into existing neural circuits (Schmidt, Strähle, and Scholpp 2013). The spinal cord, forebrain, midbrain, and hindbrain are among the primary structural components that the zebrafish brain shares with the human brain despite the zebrafish brain’s lower level of complexity (Kozol et al. 2016). These characteristics make zebrafish an excellent model for neurodevelopmental research, especially when combined with the abundance of embryos and sophisticated genetic methods for gene editing (Vaz, Hofmeister, and Lindstrand 2019). Furthermore, the transparency of zebrafish embryos allows for real-time observation of developmental processes, organogenesis, and injury responses.

Zebrafish can efficiently repair brain injuries, restoring lost function without any evident scar formation, which contrasts with the limited regenerative capabilities observed in mammals. Compared to the human brain, with 100 billion neurons, the zebrafish brain’s manageable size of approximately 100,000 neurons also makes it easier to do an in-depth study on neural growth and function (Herculano-Houzel 2009). The ease of use of automated tracking systems in high-throughput trials to evaluate diverse behaviors is made possible by this simplicity, which offers significant insights into the neurological mechanisms that underlie numerous activities like social interactions, learning, and locomotion (Kalueff, Stewart, and Gerlai 2014). Additionally, their ability to regenerate injured neural tissue, including both neurons and glia, provides valuable insights into the cellular and molecular mechanisms governing the process of regeneration. According to (X. Yu and Li 2011), zebrafish are useful in the research of ischemia brain damage and drug screening because they are easily driven into hypoxia, can have it restored, and can have medications provided directly to the water.

LncRNAs are a group of RNA molecules longer than 200 nucleotides that do not encode proteins. lncRNAs have been connected to several human diseases, such as cancer, neurological disorders, and metabolic disorders. They also play critical roles in regulating developmental processes, such as embryogenesis and tissue differentiation (Bhattacharyya et al. 2021; Aliperti, Skonieczna, and Cerase 2021). LncRNAs also play a critical role in neural cell activity during brain development and have been linked to several human brain illnesses (Andersen and Lim 2018).

LncRNAs are essential modulators of gene expression in the brain in neurodevelopmental and mental disorders and could be candidates for therapeutic intervention (Aliperti, Skonieczna, and Cerase 2021). Dysregulated long noncoding RNAs (lncRNAs) have been associated with disorders like autism, schizophrenia, and bipolar disorder, indicating their importance in maintaining normal brain development and physiology (Andersen and Lim 2018). Furthermore, lncRNAs are important in acute brain injuries such as ischemic stroke (IS). They are specifically involved in regulating gene expression and cellular processes that are essential for the brain’s repair response in ischemia-reperfusion injury (IR injury) (Wolska et al. 2021). Specific lncRNAs, including BACE1-AS, NEAT1, MALAT1, and SNHG1, which are linked to Alzheimer’s disease and may function as biomarkers or therapeutic targets, have also been linked to neurodegenerative illnesses (Asadi et al. 2021).

The purpose of this work is to discover novel lncRNAs that are expressed in the injured zebrafish brain and to understand their roles. We aim to investigate their roles in brain regeneration, inflammation, and cellular repair mechanisms by defining their expression patterns and regulatory pathways. Understanding these mechanisms could lead to novel therapeutic approaches in neuroregenerative medicine, which could have wider consequences for managing neurodegenerative illnesses and brain injuries in people.

## Methods

### Data acquisition and sample description

RNA sequencing data were downloaded from the BioStudies database (Sarkans et al. 2018) under the accession number E-MTAB-11163. The data were part of a study performed by (Demirci et al. 2022), which investigated the gene expression patterns in the telencephalon region of the zebrafish brain following Traumatic Brain Injury (TBI). The dataset comprises 18 samples, categorized into four groups: control (5 samples), 1-day post-lesion (5 samples): early wound healing stage, 3 days post-lesion (4 samples): cell proliferation stage, and 14 days post-lesion (4 samples): differentiation stage (Figure 2).

### Transcriptome Assembly and Mapping

Fastq files were downloaded from the biostudies database with Accession ID E-MTAB-11163. Raw reads were cleaned using Trimmomatic (v0.38) (Bolger, Lohse, and Usadel 2014) to remove adapters and low-quality bases using default parameters. The cleaned reads were then mapped to the zebrafish reference genome (GRCz11) using HISAT2 (v2.2.1) (Kim, Langmead, and Salzberg 2015) by disabling template length adjustment for RNA-seq reads. StringTie (v2.2.1) (Pertea et al. 2015) was used for reference-guided transcript assembly with parameters like reverse stranded library and leveraging HISAT2’s output to construct transcriptomes for each sample, followed by merging all samples (Figure 1).

**Figure 1:**
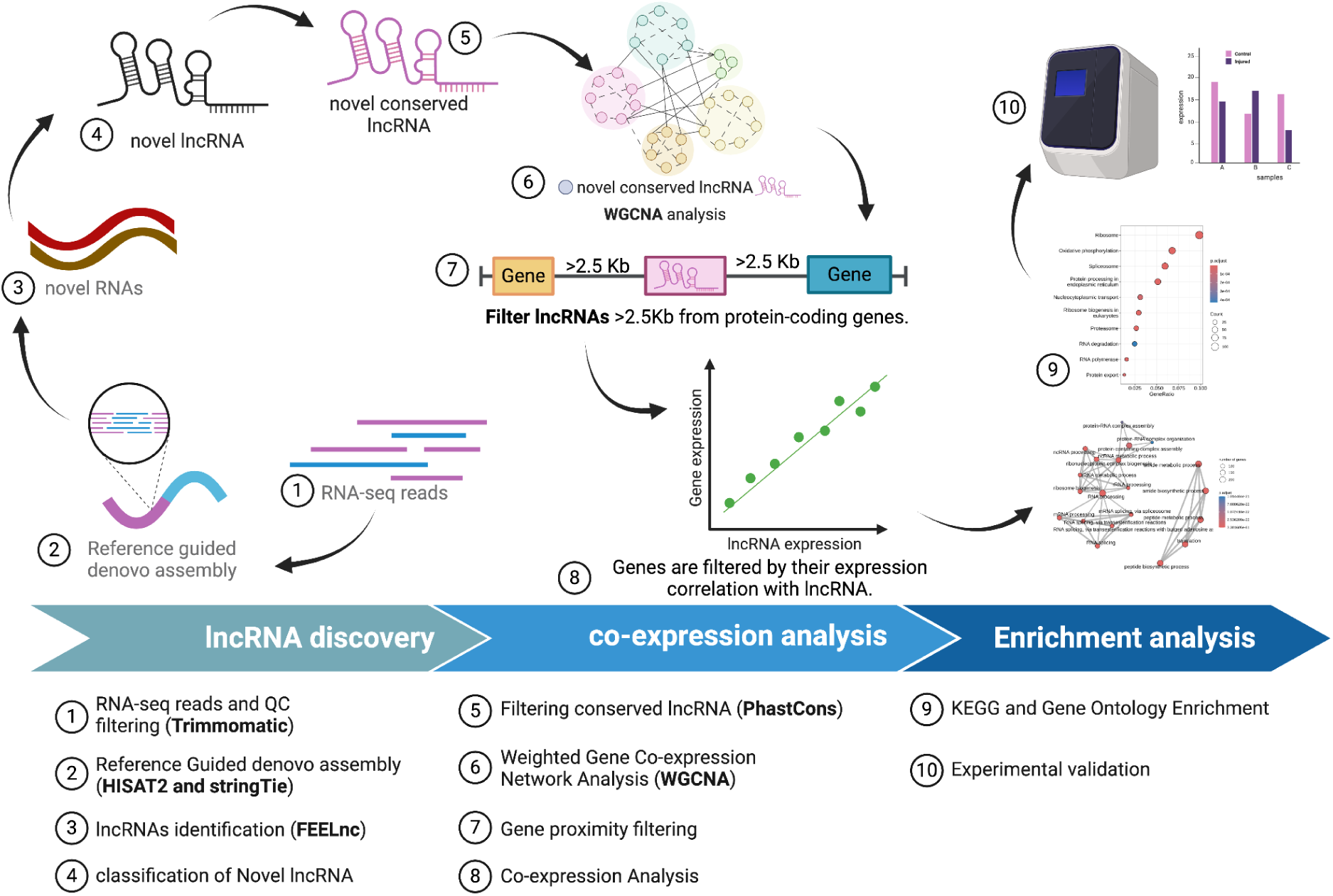
Schematic of lncRNA discovery pipeline.

### Identification and Classification of lncRNAs

To annotate and classify lncRNAs from our RNA-seq data, we employed FEELnc (FlExible Extraction of LncRNAs) (v0.2.1) (Wucher et al. 2017). This alignment-free program differentiates mRNAs from lncRNAs using a Random Forest model. This model is trained on features such as multi k-mer frequencies and relaxed open reading frames (ORFs). FEELnc has been rigorously benchmarked against leading tools and demonstrates high performance, enabling precise and reliable lncRNA identification and classification in our study.

Using Feelnc, we filtered the spurious transcripts and exons, including mono exonic regions, using FEELnc_filter.pl. After filtering, we removed the transcripts with coding potential using FEELnc_codpot.pl. Then, the lncRNA was classified based on its proximity to the annotated genes in the genome.

### Conservation Analysis

The conservation of identified lncRNA candidates across multiple species was assessed using PhastCons (V1.6) (Siepel and Haussler 2004), which employs a phylogenetic hidden Markov model to compute conservation scores. Due to the unavailability of Multiple Sequence Alignment (MSA) data for the new genome, the transcript data was first converted to version 7 of the zebrafish genome using BedFile Liftover.

### Weighted Gene Co-expression Network Analysis (WGCNA)

Weighted Gene Co-expression Network Analysis was performed using the WGCNA package (1.72-5) (Langfelder and Horvath 2008) to identify clusters of co-expressed genes and lncRNAs. Gene expression data was preprocessed, and TPM normalized prior to network construction. A soft-thresholding power of 30 was selected based on the scale-free topology criterion, and a signed network type was used to emphasize positive correlations. The network was constructed with a deep split level of 2, a minimum module size of 30 genes, and a maximum block size of 4000 genes. Modules were defined with a reassign threshold of 0 and a merge cut height of 0.25, allowing closely related modules to be merged. This analysis identified key gene networks potentially involved in the biological processes of interest.

### Gene Proximity Analysis

lncRNAs present in the brown and turquoise modules of WGCNA were further shortlisted based on their genomic location. The gene proximity analysis was performed in Python v3.11.0. Known gene coordinates were extracted from the GTF file (GRCz11.109.gtf), and lncRNA coordinates were extracted from the Stringtie results. We checked for any known gene or transcript within 2.5 kb upstream or downstream for each lncRNA. If present, that lncRNA was not considered.

The modules identified by WGCNA were subjected to further co-expression analysis to validate and refine the associations between lncRNAs and coding genes based on the presumption that lncRNAs and coding genes with similar expression patterns likely have similar functions. This analysis utilized pearson statistical correlation tests, focusing on correlations greater than 0.9 to ensure the robustness and biological significance of these relationships. Special attention was given to lncRNAs that consistently appeared central within the significant modules.

### Pathway Enrichment Analysis

Pathway enrichment analysis was conducted using the ClusterProfiler (G. Yu et al. 2012) R package to explore the potential biological functions and pathways associated with genes co-expressed with novel lncRNA clusters. This analysis helped identify enriched biological themes and pathways that may be crucially regulated by or associated with the identified novel lncRNAs, thus providing insights into their functional roles in zebrafish development and physiology. Gene ontology enrichment of biological processes and KEGG pathway enrichment are performed with a P-value less than 0.05, and all the expressed genes are chosen as background for enrichment analysis.

### Zebrafish maintenance and induction of Stab wound injury

The zebrafish were maintained at 25–28°C with a set photoperiod of 14 hours of light and 10 hours of darkness. Five to eight-month-old fish were utilized for the experiment. To investigate brain regeneration in zebrafish, a telencephalic stab wound injury was induced, and the zebrafish were momentarily anesthetized with MS222 (ethyl 3-aminobenzoate methanesulfonic acid). A 30G needle was inserted perpendicularly to the skull on the right hemisphere of the telencephalon to cause a telencephalic stab-wound injury. Thereafter, they were returned to the system water to heal. When evaluating the regeneration activity, the contralateral left hemisphere serves as the control.

### RNA isolation, cDNA synthesis, and qRT-PCR

The regenerating brains at 1dpl (day post-lesion) were harvested, and the injured telencephalon were collected (5 pooled per sample). The uninjured telencephalon (5 pooled per sample) from the same brain were collected as control. RNA isolation was done using the RNeasy Lipid Tissue Mini Kit (Cat: 1023539) according to the manufacturer’s protocol. An equal amount of RNA was used to synthesize cDNA using Takara PrimeScript 1st strand cDNA Synthesis Kit (Cat: 6110A) according to the manufacturer’s protocol. An equal amount of cDNA was used for qRT-PCR using gene-specific primers (*MSTR 23642.1, MSTR 39546.3, vegaa, nsfl1c, rac3,* and *mdm4*) and *gapdh* (F-GATACACGGAGCACCAGGTT, R-GCCATCAGGTCACATACACG) as housekeeping gene using iTaq Universal SYBR Green Supermix (Biorad, Cat: 1725121) on a Thermo Cycler (BIO-RAD CFX96). Two biological replicates and two technical replicates of each sample were used, the data was analyzed using ΔΔ Ct method, and the fold change was calculated using the 2^-ΔΔCt^ method.

## Results

The RNA sequencing retrieved from the BioStudies database consisted of 18 samples. These samples were categorized based on the timeline of post-traumatic brain injury (TBI) in zebrafish, an approach inspired by (Demirci et al. 2022). The samples included control (5 samples), 1 day post-lesion (5 samples), 3 days post-lesion (4 samples), and 14 days post-lesion (4 samples) (Figure 2). This classification was instrumental in investigating the dynamic changes in gene expression associated with different stages of brain regeneration.

**Figure 2:**
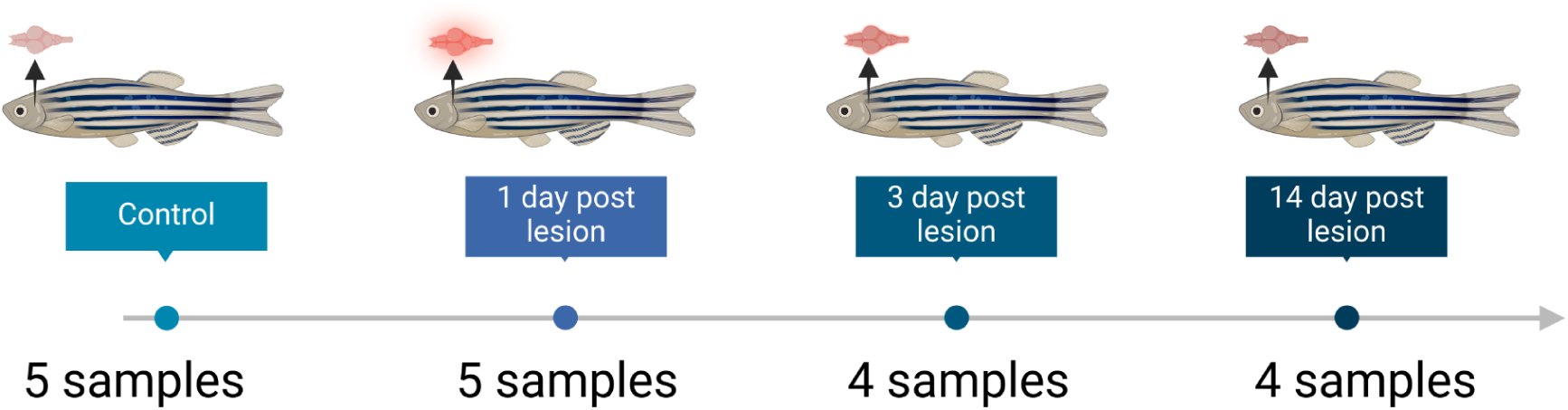
Schematic diagram illustrating the experimental design for assessing gene expression changes post-brain injury in zebrafish. The study included four groups: control, 1 day post-lesion (initial wound healing), 3 days post-lesion (cell proliferation), and 14 days post-lesion (differentiation). Each time point post-lesion represents a critical stage in the brain’s regeneration process.

### Transcriptome Assembly and Mapping

The initial quality assessment of our raw sequencing data, conducted using FastQC, demonstrated high-quality metrics across all samples, with an average Phred score of 35. After using Trimmomatic to remove adapters and low-quality bases, we aligned the cleaned data to the zebrafish reference genome (GRCz11) via HISAT2. This alignment achieved a high mapping efficiency, with an average of 88% of reads aligning successfully. Transcriptome assembly with StringTie integrated these alignments, identifying roughly 119,757 transcripts, including known and novel isoforms. We excluded transcripts derived from unplaced zebrafish genome scaffolds to enhance data relevance, focusing only on those from defined chromosomes. This refinement reduced the number of considered transcripts to 100,408.

### Identification of potential LncRNA candidates

We utilized the FEELnc tool to categorize and analyze the transcripts from our transcriptome assembly, classifying them into long non-coding RNAs (lncRNAs) and mRNAs. The FEELnc analysis successfully identified 11,281 transcripts as potential lncRNAs, characterized by their lack of coding potential, specific multi-k-mer frequencies, and open reading frame (ORF) characteristics.

To assess the evolutionary significance of these lncRNAs, we utilized the PhastCons tool, which applies a phylogenetic hidden Markov model across various vertebrate genomes. Our analysis showed that while most lncRNAs displayed low or zero conservation, mRNAs demonstrated higher conservation scores (Supplementary Figure 1). This aligns with the generally accepted understanding that lncRNA exhibits low conservation. Given the typically low conservation of lncRNAs, we hypothesized that those with any degree of conservation might play significant biological roles. Consequently, we focused on lncRNAs that exhibited non-zero conservation scores, narrowing our field from 11,281 to 6,600 transcripts. We selected the top 10% from this subset based on conservation scores, identifying 689 lncRNAs (Supplementary Figure 1, Supplementary Table 1) with notably high conservation (over 88%). This select group of lncRNAs, due to their substantial conservation, suggests potential evolutionary importance and functional relevance, making them prime candidates for further in-depth functional analyses.

**Supplementary Figure 1:**
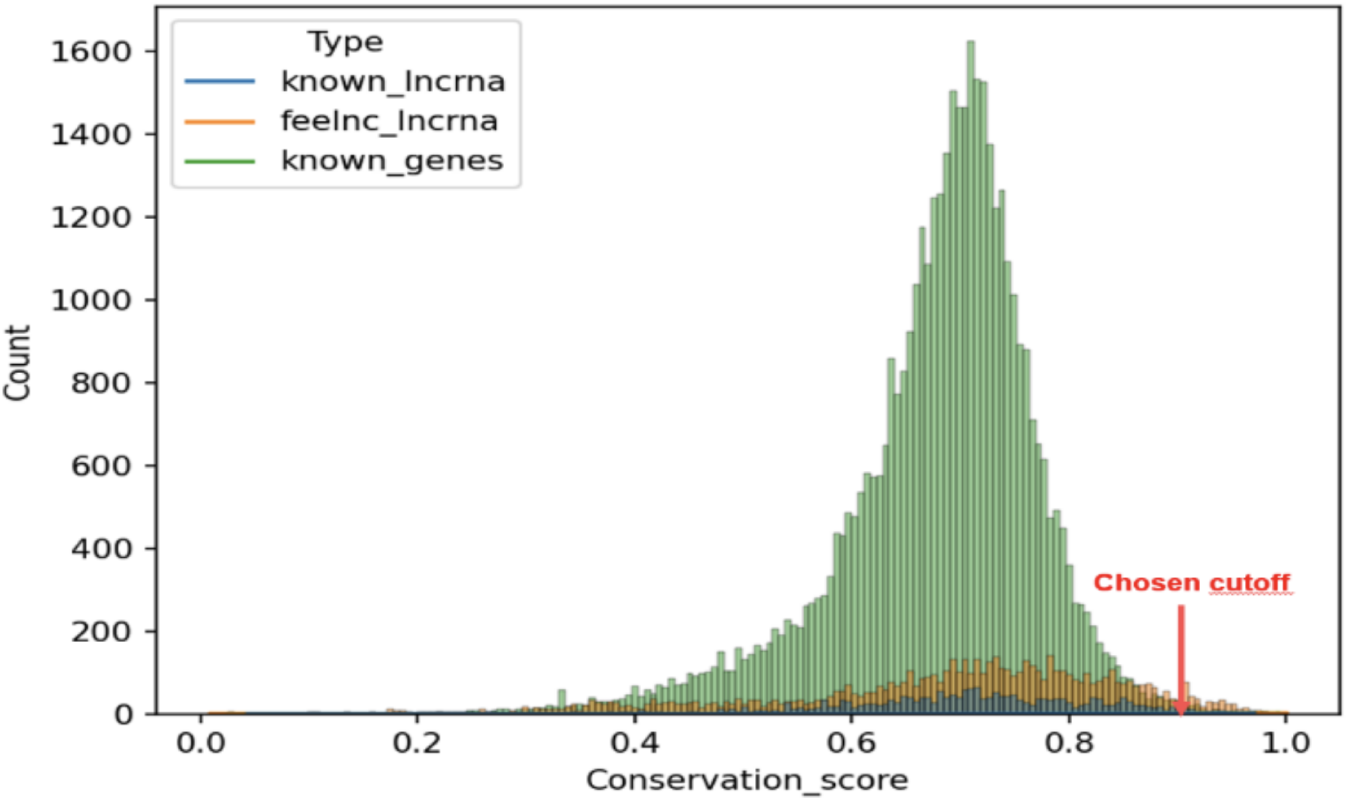
PhastCons Score distribution for various transcripts.

We assessed the expression profiles of identified lncRNA candidates across various samples using principal component analysis (PCA) of normalized lncRNA counts. The PCA results showed that samples from 1 day post-lesion (1dpl) distinctly clustered apart from other stages, which were closer to control samples (Figure 3A). This shows that the novel lncRNAs are highly active immediately after injury and slowly diminish during recovery. This pattern was corroborated by a sample-to-sample distance heatmap (Supplementary Figure 2), where 1dpl samples formed a separate cluster. These observations suggest that the expression of these lncRNAs is particularly altered during the initial stage of wound healing, possibly playing critical roles in the initial stage of the brain regeneration process.

**Figure 3:**
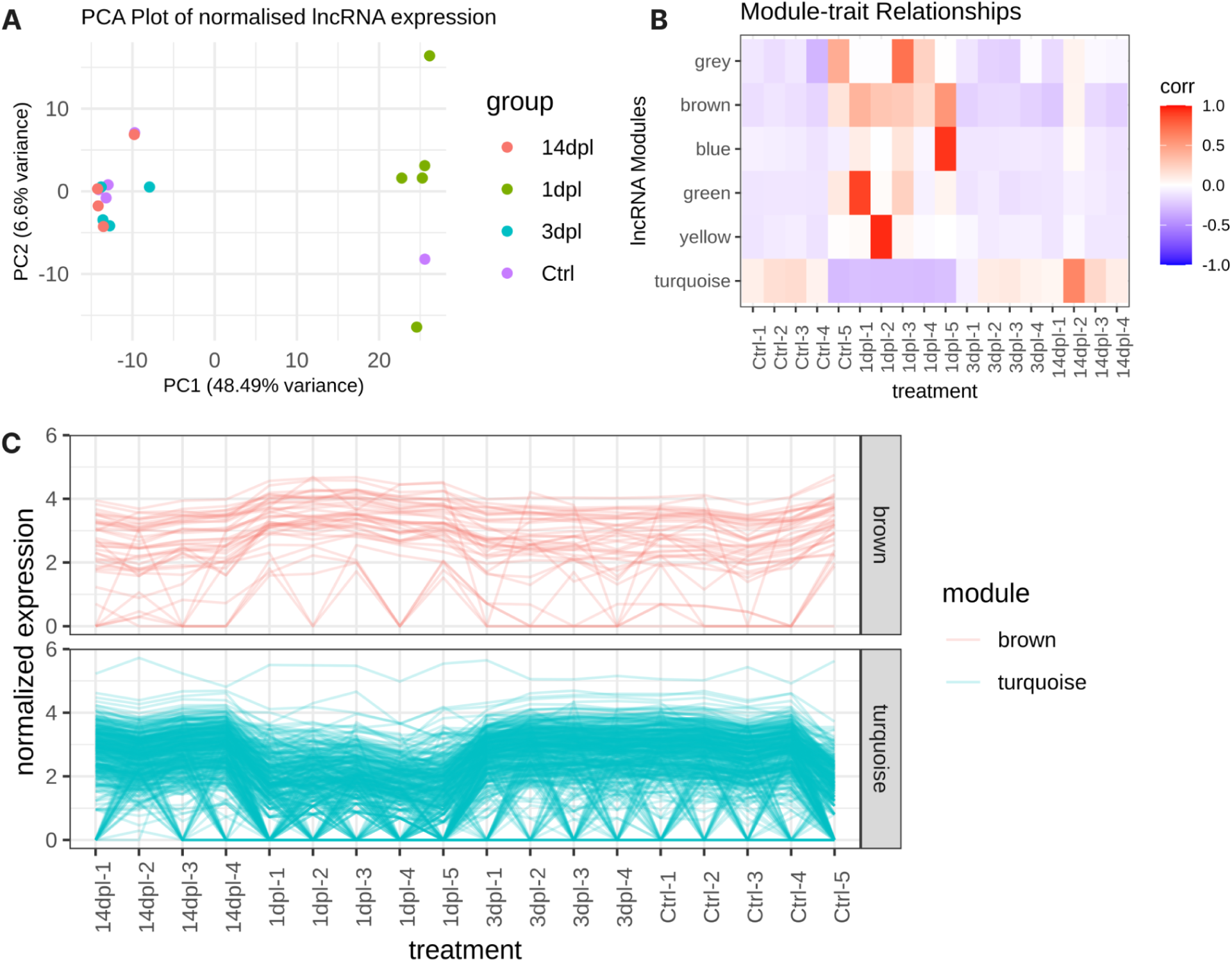
PCA and weighted gene co-expression network analysis (WGCNA) **A.** PCA plot of normalized lncRNA expression. **B.** Heatmap showing the correlation of expression profiles among lncRNA clusters. Brown and turquoise modules show consistently opposite expression profiles. **C.** Expression profiles of the individual long non-coding RNAs in the brown and turquoise co-expressed cluster. The Y-axis indicates normalized gene expression, and the X-axis indicates samples labeled according to the treatment.

**Supplementary Figure 2:**
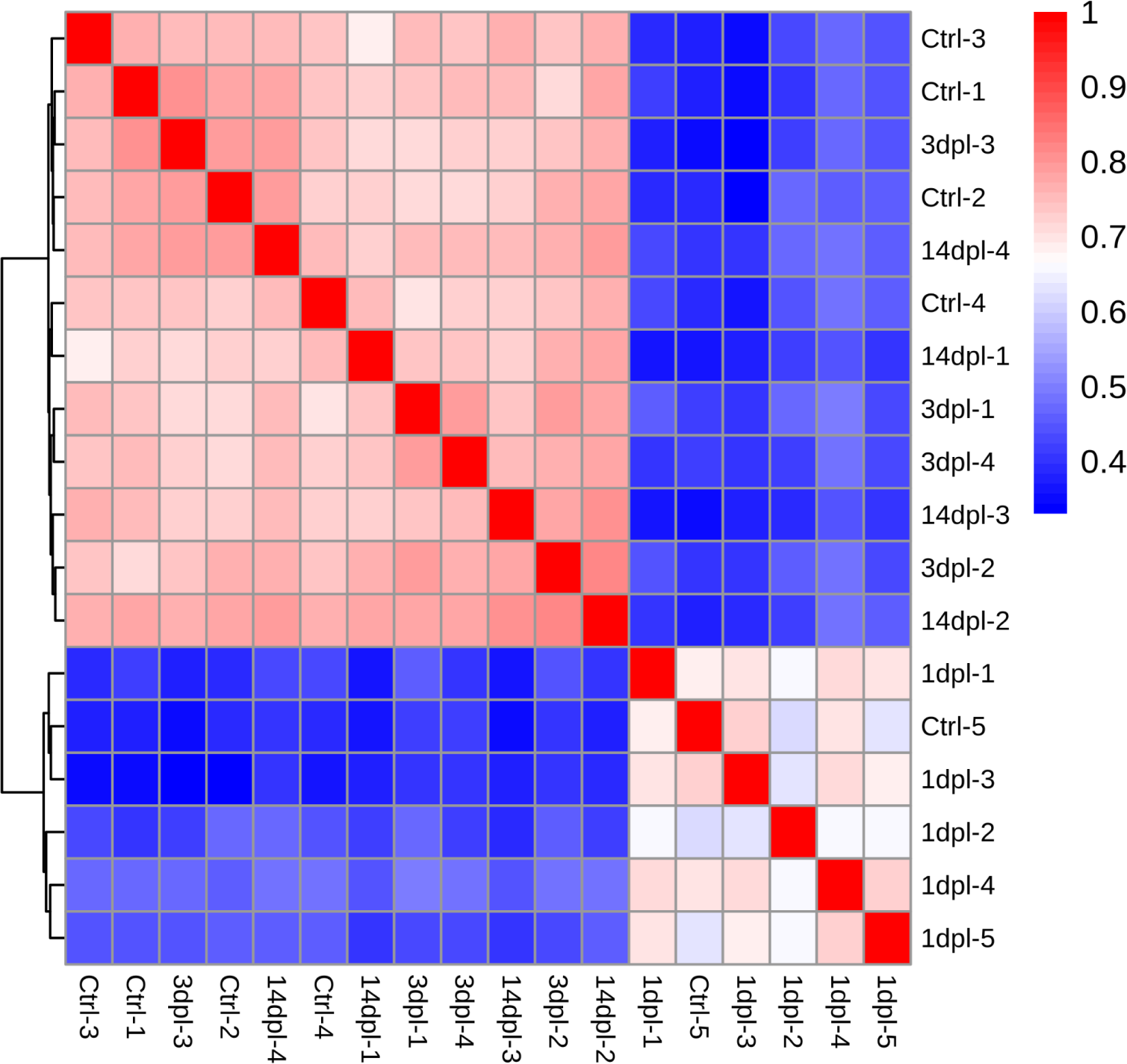
Heatmap of lncRNA expression correlation between samples. The expression profiles of one-day post-lesion samples differ from those of other samples.

We performed Weighted Gene Co-expression Network Analysis (WGCNA) to pinpoint clusters of highly coexpressed novel lncRNAs with distinct sample-specific expression patterns. This approach aimed to identify novel lncRNAs that may regulate brain regeneration. Our analysis identified six modules (Supplementary Figure 2) with unique expression profiles across the brain injury recovery timeline (Figure 3B), with particular attention to two modules, brown and turquoise, that demonstrated unique and consistent patterns at the 1-day post-lesion (1dpl) stage. The brown module showed increased expression, while the turquoise module displayed decreased expression at 1dpl (Figure 3C). We identified 385 novel lncRNAs in the turquoise module and 39 in the brown module (Supplementary Figure 3, Supplementary Table 1). To ensure the accuracy of our functional inference, we implemented stringent criteria, excluding any novel lncRNAs located within 2.5 kb of coding genes to avoid potential interference. This filtration left us with 77 lncRNAs in the turquoise module and 9 in the brown module (Table 1, Supplementary Table 2), isolating those with potentially independent functional roles.

**Table 1:** Gene proximity analysis filtering table.

| Module | Before proximity<br>Filtering | After proximity<br>Filtering |
| --- | --- | --- |
| Turquoise | 385 | 77 |
| Brown | 39 | 9 |

**Supplementary Figure 3:**
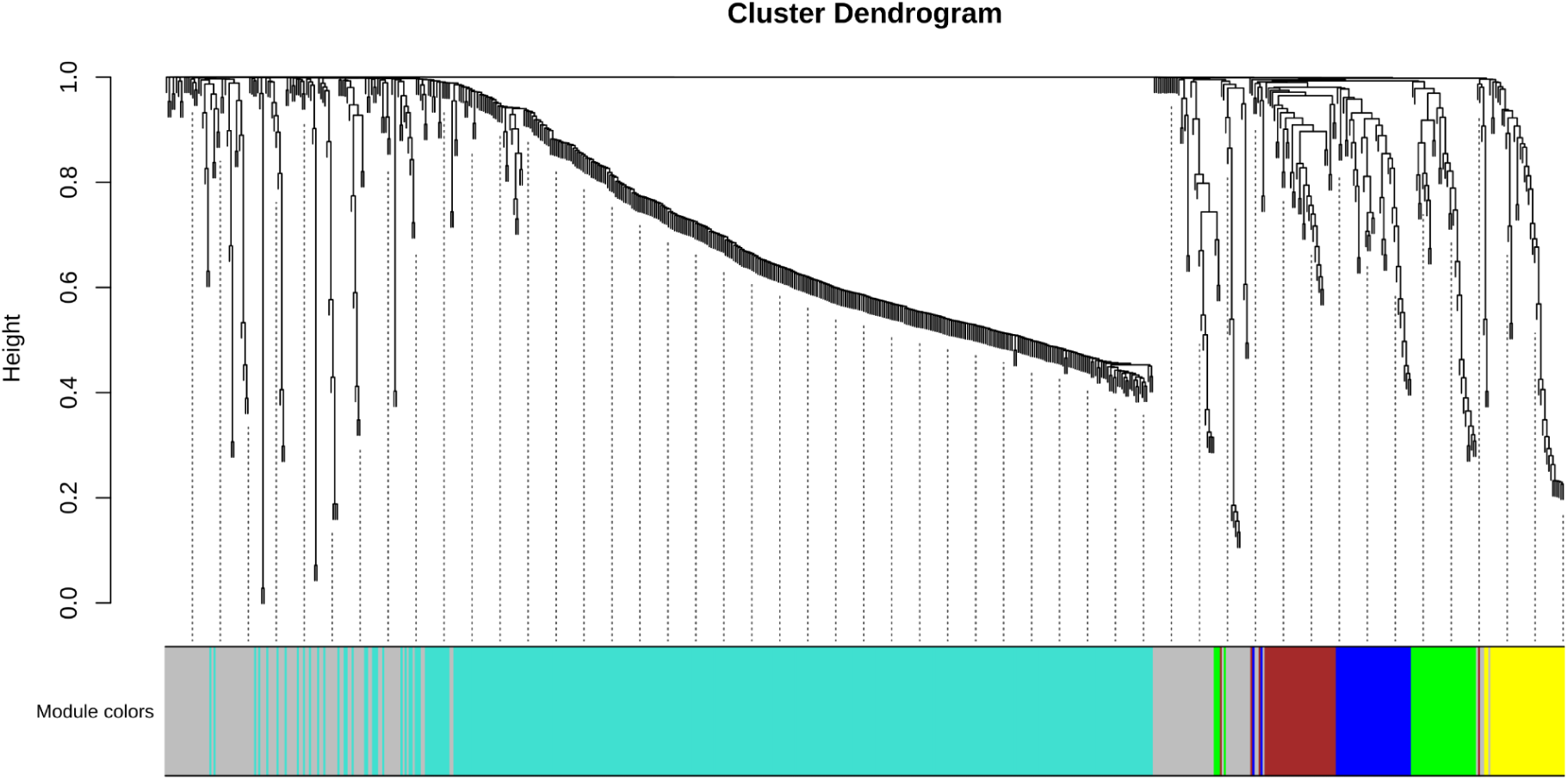
Dendrogram of WGCNA Clusters with 689 novel lncRNA.

### Functional significance of identified lncRNAs

To further explore the biological significance of these identified lncRNAs, we extended our analysis by correlating these novel lncRNAs with expressed known genes. We filtered genes that showed a high correlation (cutoff > |0.9|) with the novel lncRNAs and conducted gene ontology (GO) and KEGG pathway enrichment analyses, aiming to elucidate the biological processes and pathways potentially influenced by these lncRNAs during the initial wound healing stage of brain regeneration.

Genes correlating to brown module lncRNAs show enrichment for metabolic processes related to RNA splicing and peptide biosynthetic processes (Figure 7), suggesting the role of these lncRNAs in these processes.

GO terms enriched for the brown module include peptide metabolic process, amide metabolic process, and translation (Figure 4A) while the KEGG pathway enrichment analysis of brown module highlight significant enrichment in pathways related to the ribosome, oxidative phosphorylation, and the spliceosome (Figure 4B). This suggests that genes involved in protein synthesis, peptide metabolism, energy production, and RNA splicing are highly represented in brown module gene set during early wound healing stages. This aligns with known associations of metabolic distress, peptide mechanisms and splicing with brain injury (Gourain et al. 2021; Lakshmanan et al. 2010; Ilieş, Zupanc, and Zupanc 2012), suggesting that lncRNAs of brown module could potentially play a role in regulating these processes in response to brain injury and recovery.

**Figure 4:**
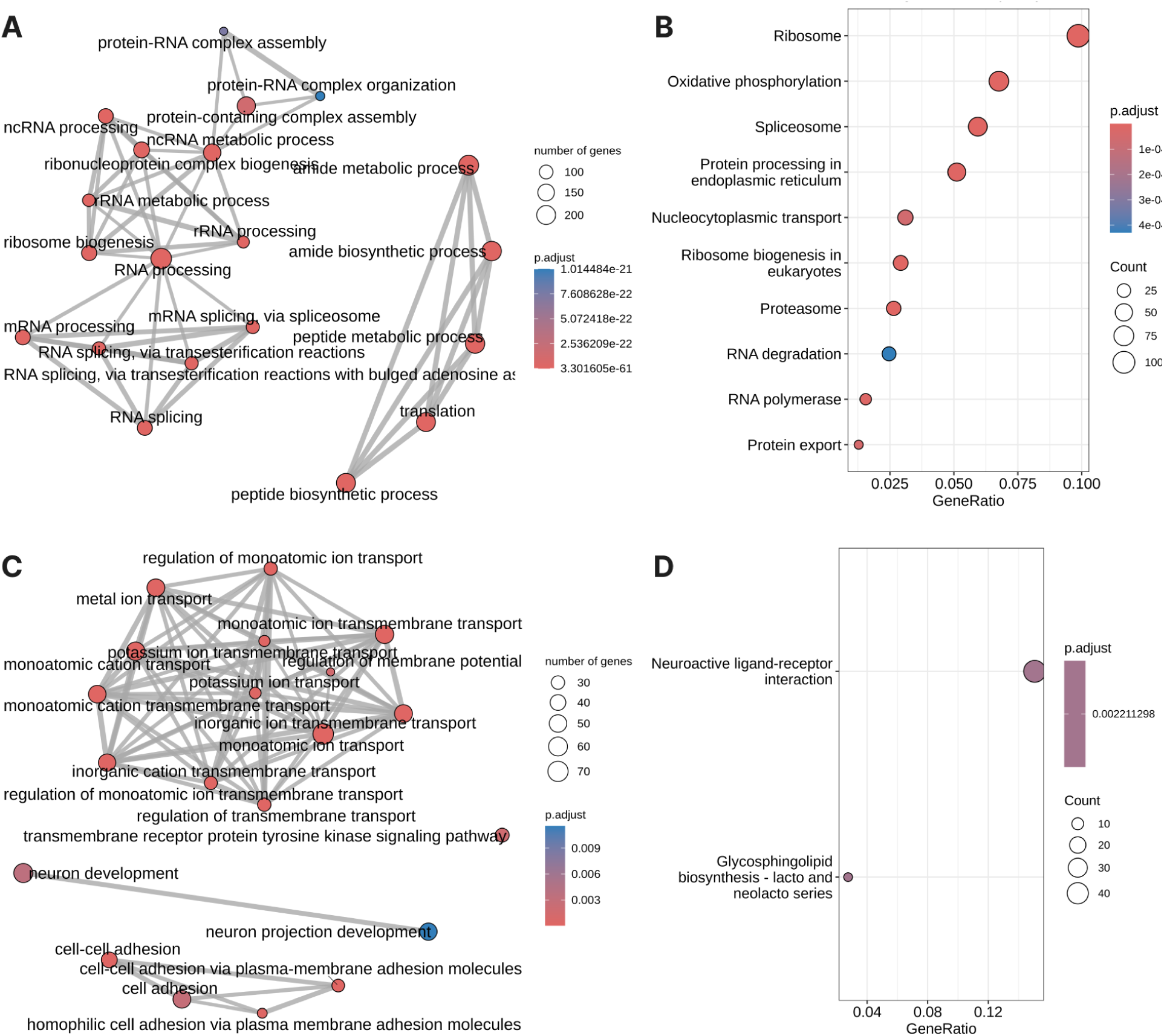
Functional Enrichment Analysis. **A.** EMAP plot showing GO enrichment of biological processes for genes with a 90% correlation in expression with lncRNAs in the brown module. **B.** Dot plot of KEGG pathway enrichment analysis for genes highly correlated (r > 0.90) with lncRNAs in the brown module. **C.** EMAP plot showing GO enrichment of biological processes for genes with a 90% correlation in expression with lncRNAs in the turquoise module. **D.** Dot plot of KEGG pathway enrichment analysis for genes highly correlated (r > 0.90) with lncRNAs in the turquoise module

In the turquoise module, novel lncRNAs are predominantly linked to regulation of ion transmembrane transport, metal ion transmembrane transport, and potassium ion transmembrane transport. These processes are critical for maintaining cellular homeostasis and membrane potential, which are essential for neuron firing and muscle contraction (Figure 4C). Furthermore, KEGG pathway analysis of the gene set correlated to turquoise module lncRNAs highlights significant enrichment in the neuroactive ligand-receptor interaction pathway and the glycosphingolipid biosynthesis - lacto and neolacto series pathway (Figure 4D). These pathways play vital roles in neuronal signaling, cell-cell communication, and membrane integrity (D’Angelo et al. 2013; J. Wei et al. 2020), suggesting that lncRNAs in this module might regulate these critical processes.

Interestingly, the neuroactive ligand-receptor interaction pathway is well-documented for its significant role in various cancers (Subramanian and Devarajan 2022; P. Wei, Tang, and Li 2012; Huan et al. 2014). Demicri and colleagues also showed that differentially expressed genes at one day post-lesion (1dpl) share a maximum number of genes with brain cancer, compared to 3dpl and 14dpl (Demirci et al. 2022). This suggests a potential regulatory role of lncRNAs in the neuroactive ligand-receptor interaction pathway, not only in maintaining neuronal function but also possibly in pathological states such as cancer.

**Figure 5:**
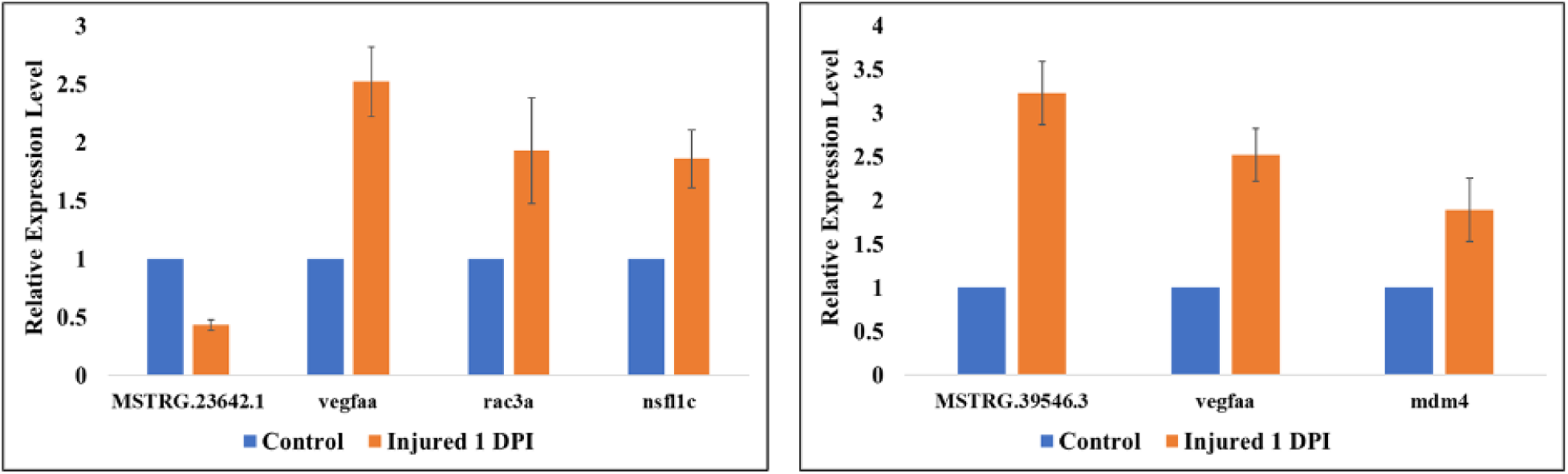
Relative lncRNA and gene expression level in the pooled five hemispheres of telencephalon during the early wound injury at 1 dpi. Error bars represent ± standard error of the mean (SEM, n = 2). (A) MSTR 23642.1 lncRNA expression in combination with vegaa, rac3a, and nsfl1c genes and (B) MSTR 39546.3 lncRNA expression in combination with vagaa and mdm4 genes.

We chose 2 lncRNAs to validate this study experimentally, one from brown (MSTRG 395546.3) and another from turquoise (MSTRG 23642.1) modules and chose specific gene pairs to perform experimental validation of co-expression using quantitative PCR. Since brown and turquoise show opposite expression pattern in 1-dpl we chose gene pairs having positive correlation with lncRNA from brown module and negatively correlated genes with lncRNA of turquoise module. qPCR validation of lncRNA and gene pairs were performed. qPCR confirmed the in silico results that lncRNA in the MSTRG 23642 (brown module) positively correlate with genes vegfaa, rac3a and nsfl1c and genes vegfaa and mdm4 negatively correlate with lncRNA MSTRG 39546.3 (turquoise module).

The vascular endothelial growth factor Aa (vegfaa) gene is central to the angiogenesis process (Marín-Juez et al. 2016). Angiogenesis, activated within a few days post-cerebral ischemia, plays a crucial role in neuronal remodeling and functional recovery. It provides guidance to sprouting axons and enhances the proliferation and differentiation of neural stem cells (Wang et al. 2007; Kanazawa et al. 2017; Ruan et al. 2015; Hatakeyama, Ninomiya, and Kanazawa 2020). Demirci et al. 2022 also observed an upregulation of vegfaa specifically at 1 day post-lesion (1 dpl), suggesting that injury triggers rapid angiogenic sprouting during early brain regeneration (Demirci et al. 2022). The negative correlation between lncRNAs in the turquoise module and vegfaa indicates that these lncRNAs might repress or inhibit vegfaa expression. The downregulation of these lncRNAs at 1 dpl may relieve their inhibitory effect on vegfaa, allowing for the necessary upregulation of vegfaa to support angiogenesis and neural regeneration processes. Additionally, the gene rac3a, associated with heart regeneration pathways, is also negatively correlated with turquoise module lncRNAs MSTRG 23642.1.

On the other hand, The positive correlation between lncRNAs in the brown module and vegfaa suggests that these lncRNAs might support the upregulation of vegfaa, promoting angiogenesis and facilitating the initial stages of brain regeneration. These lncRNAs could be acting as enhancers or stabilizers of vegfaa expression, thereby contributing to the rapid angiogenic response observed at 1 dpl. Mdm4 also shows a positive correlation with brown lncRNA MSTRG 23642.1. The MDM4/MDM2-p53-IGF1 axis, which controls axonal regeneration, sprouting, and functional recovery after CNS injury (Joshi et al. 2015), shows a positive correlation with lncRNAs in the brown module, suggesting that these lncRNAs may play a role in these critical regenerative processes as well.

Thus, our findings suggest a dual regulatory mechanism where lncRNAs in the brown module promote the expression of genes participating in brain regeneration processes, while lncRNAs in the turquoise module inhibit genes involved in the regeneration and recovery of the brain.

## Discussion

Building on a recent study by Demirci et al., 2022, on gene expression during brain regeneration post-injury in zebrafish, our research investigated the regulatory role of long non-coding RNAs(lncRNAs) in this process. Demirci et al.’s work provided crucial insights into the dynamic gene expression changes across three key stages of brain regeneration. They discovered that the most significant changes occurred in the initial wound healing stage (one-day post-lesion), with notable similarities between these expression patterns and those observed in brain cancer.

We utilized their published RNA-seq data to identify and characterize lncRNAs that might influence these expression changes. The study of lncRNAs is vital as these molecules play key roles in regulating gene expression, and their dysregulation has been implicated in various pathologies, including cancer and potentially in the mechanisms of neuronal repair and regeneration (Hezroni, Perry, and Ulitsky 2019; Jiang et al. 2019).

The raw RNA seq was first refined to ensure quality and accuracy, followed by alignment using HISAT2 and transcriptome assembly using StringTie. Out of the 100,408 transcripts assembled by StringTie, FEELnc identified 11,281 potential lncRNAs based on coding potential and other structural features. To further refine our selection, we applied PhastCons conservation scores, which reduced the number of potential lncRNAs to 689 transcripts. Given that lncRNAs typically exhibit low conservation (Johnsson et al. 2014) across species, we prioritized those with high PhastCons scores to ensure the selection of functionally relevant lncRNAs.

The PCA plot and heatmap analysis of the 689 lncRNA candidates distinctly demonstrated that samples from the 1-day post-lesion (1dpl) stage clustered separately from other stages. Interestingly, while Demirci et al. showed distinct gene expression profiles at both the 1-day (initial wound healing) and 3-day (cell proliferation) stages, with the 14-day profile mostly resembling controls while our study observed that lncRNA expression profiles exhibited a distinct pattern primarily at the 1-day post-injury stage. This suggests that the expression patterns of identified lncRNAs are markedly distinct during the initial stage of wound healing, potentially playing important roles in initial wound healing roles. The significant divergence in lncRNA expression specifically at 1dpl underscores their possible involvement in early regulatory mechanisms critical to initiating and guiding the regenerative processes.

To ascertain the biological functions of these lncRNA candidates, we performed a WGCNA analysis. WGCNA revealed six modules with distinct expression profiles over the brain injury recovery timeline. Notably, two modules (brown and turquoise) exhibited consistent unique expression patterns specific to the 1-day post-lesion (1dpl) stage, with the brown module showing increased expression and the turquoise module showing decreased expression at this early stage. This pattern aligns with our previous observations of significant lncRNA activity during the initial wound-healing phase. We extended our analysis by correlating these lncRNAs with known genes and conducted gene ontology (GO) and pathway enrichment analysis of genes highly correlated (cutoff > |0.9|) with these lncRNAs. The analysis revealed that the peptide biosynthetic process, cellular amide metabolic process, and cellular macromolecule biosynthetic process are interconnected pathways involved in producing the building blocks of life.

Further, the GO analysis highlighted the critical roles of lncRNAs in the response to traumatic brain injury. The genes within the brown module were enriched in functions related to peptide and amide metabolism, translation; and pathways involving the ribosome, oxidative phosphorylation, and the spliceosome. These pathways are associated with protein synthesis, energy production, and RNA splicing, indicating the important role of lncRNA in regulating genes involved in these processes during early wound healing (Gourain et al. 2021; Lakshmanan et al. 2010; Ilieş, Zupanc, and Zupanc 2012).

In the turquoise module, lncRNAs are associated with ion transmembrane transport, crucial for cellular homeostasis and membrane potential necessary for neuron firing and muscle contraction. KEGG pathway analysis shows enrichment in neuroactive ligand-receptor interaction and glycosphingolipid biosynthesis pathways, important for neuronal signaling, cell communication, and membrane integrity (D’Angelo et al. 2013; J. Wei et al. 2020).

Moreover, we selected two lncRNA targets and their highly correlated gene pairs, showing both negative and positive correlations, for experimental validation using qPCR. The qPCR results confirmed the expression patterns observed in our RNA-seq analysis, providing further support for the regulatory role of these lncRNAs in brain regeneration. These experimentally validated lncRNAs and their correlated genes suggest specific regulatory mechanisms where lncRNAs might act as key modulators of gene expression, either enhancing or suppressing their correlated targets during the early stages of wound healing.

Our findings suggest that specific lncRNAs are intricately involved in the early stages of brain regeneration in zebrafish, potentially regulating key genes and pathways that drive the regenerative process. These insights not only enhance our understanding of lncRNA function in neuronal repair but also provide a foundation for future research aimed at leveraging lncRNAs as therapeutic targets for brain.

## Data availability

Sequencing data files can be downloaded from the biostudies database with Accession ID E-MTAB-11163. The bioinformatic analysis pipeline will be provided upon request to the corresponding authors.

## Supplementary Files

Supplementary Table 1: List of lncRNA gene IDs and their respective module in WCGNA

Supplementary Table 2: List of lncRNA transcript IDs from brown and turquoise module

## References

Aliperti, Vincenza, Justyna Skonieczna, and Andrea Cerase. 2021. “Long Non-Coding RNA (lncRNA) Roles in Cell Biology, Neurodevelopment and Neurological Disorders.” Non-Coding RNA 7 (2). 10.3390/ncrna7020036.

Andersen, Rebecca E., and Daniel A. Lim. 2018. “Forging Our Understanding of lncRNAs in the Brain.” Cell and Tissue Research 371 (1): 55–71.

Asadi, Mohammad Reza, Mehdi Hassani, Shiva Kiani, Hani Sabaie, Marziyeh Sadat Moslehian, Mohammad Kazemi, Soudeh Ghafouri-Fard, Mohammad Taheri, and Maryam Rezazadeh. 2021. “The Perspective of Dysregulated LncRNAs in Alzheimer’s Disease: A Systematic Scoping Review.” Frontiers in Aging Neuroscience 13 (September): 709568.

Bhattacharyya, Nirjhar, Vedansh Pandey, Malini Bhattacharyya, and Abhijit Dey. 2021. “Regulatory Role of Long Non Coding RNAs (lncRNAs) in Neurological Disorders: From Novel Biomarkers to Promising Therapeutic Strategies.” Asian Journal of Pharmaceutical Sciences 16 (5): 533–50.

Bolger, Anthony M., Marc Lohse, and Bjoern Usadel. 2014. “Trimmomatic: A Flexible Trimmer for Illumina Sequence Data.” Bioinformatics 30 (15): 2114–20.

D’Angelo, Giovanni, Serena Capasso, Lucia Sticco, and Domenico Russo. 2013. “Glycosphingolipids: Synthesis and Functions.” The FEBS Journal 280 (24): 6338–53.

Demirci, Yeliz, Guillaume Heger, Esra Katkat, Irene Papatheodorou, Alvis Brazma, and Gunes Ozhan. 2022. “Brain Regeneration Resembles Brain Cancer at Its Early Wound Healing Stage and Diverges From Cancer Later at Its Proliferation and Differentiation Stages.” Frontiers in Cell and Developmental Biology 10 (February): 813314.

Gourain, Victor, Olivier Armant, Luisa Lübke, Nicolas Diotel, Sepand Rastegar, and Uwe Strähle. 2021. “Multi-Dimensional Transcriptome Analysis Reveals Modulation of Cholesterol Metabolism as Highly Integrated Response to Brain Injury.” Frontiers in Neuroscience 15 (May): 671249.

Hatakeyama, Masahiro, Itaru Ninomiya, and Masato Kanazawa. 2020. “Angiogenesis and Neuronal Remodeling after Ischemic Stroke.” Neural Regeneration Research 15 (1): 16–19.

Herculano-Houzel, Suzana. 2009. “The Human Brain in Numbers: A Linearly Scaled-up Primate Brain.” Frontiers in Human Neuroscience 3 (November): 31.

Hezroni, Hadas, Rotem Ben Tov Perry, and Igor Ulitsky. 2019. “Long Noncoding RNAs in Development and Regeneration of the Neural Lineage.” Cold Spring Harbor Symposia on Quantitative Biology 84: 165–77.

Huan, Jinliang, Lishan Wang, Li Xing, Xianju Qin, Lingbin Feng, Xiaofeng Pan, and Ling Zhu. 2014. “Insights into Significant Pathways and Gene Interaction Networks Underlying Breast Cancer Cell Line MCF-7 Treated with 17β-Estradiol (E2).” Gene 533 (1): 346–55.

Ilieş, I., M. M. Zupanc, and G. K. H. Zupanc. 2012. “Proteome Analysis Reveals Protein Candidates Involved in Early Stages of Brain Regeneration of Teleost Fish.” Neuroscience 219 (September): 302–13.

Jiang, Ming-Chun, Jiao-Jiao Ni, Wen-Yu Cui, Bo-Ya Wang, and Wei Zhuo. 2019. “Emerging Roles of lncRNA in Cancer and Therapeutic Opportunities.” American Journal of Cancer Research 9 (7): 1354–66.

Johnsson, Per, Leonard Lipovich, Dan Grandér, and Kevin V. Morris. 2014. “Evolutionary Conservation of Long Non-Coding RNAs; Sequence, Structure, Function.” Biochimica et Biophysica Acta 1840 (3): 1063–71.

Joshi, Yashashree, Marília Grando Sória, Giorgia Quadrato, Gizem Inak, Luming Zhou, Arnau Hervera, Khizr I. Rathore, et al. 2015. “The MDM4/MDM2-p53-IGF1 Axis Controls Axonal Regeneration, Sprouting and Functional Recovery after CNS Injury.” Brain: A Journal of Neurology 138 (Pt 7): 1843–62.

Kalueff, Allan V., Adam Michael Stewart, and Robert Gerlai. 2014. “Zebrafish as an Emerging Model for Studying Complex Brain Disorders.” Trends in Pharmacological Sciences 35 (2): 63–75.

Kanazawa, Masato, Minami Miura, Masafumi Toriyabe, Misaki Koyama, Masahiro Hatakeyama, Masanori Ishikawa, Takashi Nakajima, et al. 2017. “Microglia Preconditioned by Oxygen-Glucose Deprivation Promote Functional Recovery in Ischemic Rats.” Scientific Reports 7 (February): 42582.

Kim, Daehwan, Ben Langmead, and Steven L. Salzberg. 2015. “HISAT: A Fast Spliced Aligner with Low Memory Requirements.” Nature Methods 12 (4): 357–60.

Kozol, Robert A., Alexander J. Abrams, David M. James, Elena Buglo, Qing Yan, and Julia E. Dallman. 2016. “Function Over Form: Modeling Groups of Inherited Neurological Conditions in Zebrafish.” Frontiers in Molecular Neuroscience 9 (July): 55.

Lakshmanan, R., J. A. Loo, T. Drake, J. Leblanc, A. J. Ytterberg, D. L. McArthur, M. Etchepare, and P. M. Vespa. 2010. “Metabolic Crisis after Traumatic Brain Injury Is Associated with a Novel Microdialysis Proteome.” Neurocritical Care 12 (3): 324–36.

Langfelder, Peter, and Steve Horvath. 2008. “WGCNA: An R Package for Weighted Correlation Network Analysis.” BMC Bioinformatics 9 (December): 559.

Lowery, Laura Anne, Gianluca De Rienzo, Jennifer H. Gutzman, and Hazel Sive. 2009. “Characterization and Classification of Zebrafish Brain Morphology Mutants.” Anatomical Record 292 (1): 94–106.

Marín-Juez, Rubén, Michele Marass, Sebastien Gauvrit, Andrea Rossi, Shih-Lei Lai, Stefan C. Materna, Brian L. Black, and Didier Y. R. Stainier. 2016. “Fast Revascularization of the Injured Area Is Essential to Support Zebrafish Heart Regeneration.” Proceedings of the National Academy of Sciences of the United States of America 113 (40): 11237–42.

Pertea, Mihaela, Geo M. Pertea, Corina M. Antonescu, Tsung-Cheng Chang, Joshua T. Mendell, and Steven L. Salzberg. 2015. “StringTie Enables Improved Reconstruction of a Transcriptome from RNA-Seq Reads.” Nature Biotechnology 33 (3): 290–95.

Ruan, Linhui, Brian Wang, Qichuan ZhuGe, and Kunlin Jin. 2015. “Coupling of Neurogenesis and Angiogenesis after Ischemic Stroke.” Brain Research 1623 (October): 166–73.

Sarkans, Ugis, Mikhail Gostev, Awais Athar, Ehsan Behrangi, Olga Melnichuk, Ahmed Ali, Jasmine Minguet, et al. 2018. “The BioStudies Database-One Stop Shop for All Data Supporting a Life Sciences Study.” Nucleic Acids Research 46 (D1): D1266–70.

Schmidt, Rebecca, Uwe Strähle, and Steffen Scholpp. 2013. “Neurogenesis in Zebrafish - from Embryo to Adult.” Neural Development 8 (February): 3.

Siepel, Adam, and David Haussler. 2004. “Computational Identification of Evolutionarily Conserved Exons.” In Proceedings of the Eighth Annual International Conference on Research in Computational Molecular Biology, 177–86. RECOMB ’04. New York, NY, USA: Association for Computing Machinery.

Subramanian, Umadevi, and Bharanidharan Devarajan. 2022. “Identification of Dysregulated Pathways and Key Genes in Human Retinal Angiogenesis Using Microarray Metadata.” Gene Reports 26 (March): 101434.

Vaz, Raquel, Wolfgang Hofmeister, and Anna Lindstrand. 2019. “Zebrafish Models of Neurodevelopmental Disorders: Limitations and Benefits of Current Tools and Techniques.” International Journal of Molecular Sciences 20 (6). 10.3390/ijms20061296.

Wang, Yaoming, Kunlin Jin, Xiao Ou Mao, Lin Xie, Surita Banwait, Hugo H. Marti, and David A. Greenberg. 2007. “VEGF-Overexpressing Transgenic Mice Show Enhanced Post-Ischemic Neurogenesis and Neuromigration.” Journal of Neuroscience Research 85 (4): 740–47.

Wei, Jialiu, Jianhui Liu, Shuang Liang, Mengqi Sun, and Junchao Duan. 2020. “Low-Dose Exposure of Silica Nanoparticles Induces Neurotoxicity via Neuroactive Ligand-Receptor Interaction Signaling Pathway in Zebrafish Embryos.” International Journal of Nanomedicine 15 (June): 4407–15.

Wei, Peng, Hongwei Tang, and Donghui Li. 2012. “Insights into Pancreatic Cancer Etiology from Pathway Analysis of Genome-Wide Association Study Data.” PloS One 7 (10): e46887.

Wolska, Marta, Joanna Jarosz-Popek, Eva Junger, Zofia Wicik, Tahmina Porshoor, Lucia Sharif, Pamela Czajka, et al. 2021. “Long Non-Coding RNAs as Promising Therapeutic Approach in Ischemic Stroke: A Comprehensive Review.” Molecular Neurobiology 58 (4): 1664–82.

Wucher, Valentin, Fabrice Legeai, Benoît Hédan, Guillaume Rizk, Lætitia Lagoutte, Tosso Leeb, Vidhya Jagannathan, et al. 2017. “FEELnc: A Tool for Long Non-Coding RNA Annotation and Its Application to the Dog Transcriptome.” Nucleic Acids Research 45 (8): e57.

Yu, Guangchuang, Li-Gen Wang, Yanyan Han, and Qing-Yu He. 2012. “clusterProfiler: An R Package for Comparing Biological Themes among Gene Clusters.” Omics: A Journal of Integrative Biology 16 (5): 284–87.

Yu, Xinge, and Yang V. Li. 2011. “Zebrafish as an Alternative Model for Hypoxic-Ischemic Brain Damage.” International Journal of Physiology, Pathophysiology and Pharmacology 3 (2): 88–96.

